# RNA polymerase II is recruited to DNA double-strand breaks for dilncRNA transcription in *Drosophila*

**DOI:** 10.1101/2021.06.10.447683

**Authors:** Romy Böttcher, Ines Schmidts, Volker Nitschko, Petar Duric, Klaus Förstemann

## Abstract

DNA double-strand breaks are among the most toxic lesions that can occur in a genome and their faithful repair is thus of great importance. Recent findings have uncovered local transcription that initiates at the break and forms a non-coding transcript, called damage-induced long non-coding RNA (dilncRNA), which helps to coordinate the DNA transactions necessary for repair. We provide nascent RNA sequencing-based evidence that RNA polymerase II transcribes the dilncRNA in *Drosophila* and that this is more efficient for DNA breaks in an intron-containing gene, consistent with the higher damage-induced siRNA levels downstream of an intron. The spliceosome thus stimulates recruitment of RNA polymerase II to the break, rather than merely promoting the annealing of sense and antisense RNA to form the siRNA precursor. In contrast, RNA polymerase III nascent RNA libraries did not contain reads corresponding to the cleaved loci and selective inhibition of RNA polymerase III did not reduce the yield of damage-induced siRNAs. Finally, the damage-induced siRNA density was unchanged downstream of a T8 sequence, which terminates RNA polymerase III transcription. We thus found no evidence for a participation of RNA polymerase III in dilncRNA transcription in cultured Drosophila cells.

## Introduction

The siRNA silencing system in *Drosophila* helps to fend off viral infections [1], but also contributes to the control of transposon mobilization in somatic cells [2]. In both cases, the trigger for siRNA generation is double-stranded RNA (dsRNA). During viral infection, this likely stems from replication intermediates, while for genome surveillance convergent transcription must occur. For multi-copy sequences, this convergent transcription can also be envisaged to occur *in trans*, i.e. at different instances of the same sequence. A particular form of dsRNA generation has been identified in *Drosophila* at transcribed DNA double-strand breaks [3]. The genetic requirements indicate an involvement of the spliceosome and this appears to be true for the surveillance of high-copy sequences as well [4]. Intriguingly, stalled spliceosomes can recruit RNA-dependent RNA polymerase (RdRP) to transposon mRNAs in the pathogenic yeast *Cryptococcus neoformans* [5]. For organisms that lack an RdRP gene, however, induction of convergent transcription must happen at the DNA.

DNA double-strand breaks (DSB) are highly toxic genome lesions that need to be faithfully repaired. A series of molecular interactions is initiated once a DSB has been detected and signaling events recruit repair factors, modify local chromatin structure and mitigate access between transcription and DNA repair proteins [6]. Many studies have concluded that a relatively large region around the DSB is transcriptionally silenced in a reversible manner, presumably to avoid conflicts between transcription and repair [7]. In recent years, however, antisense transcription that initiates at the DNA break has been observed [8-10]. In the context of DNA repair, this transcription seems to fine-tune the dose of single-strand binding proteins such as RPA that initially associate with the 3’->5’ resected break [8]. Furthermore, damage-induced small RNAs derived from these antisense transcripts have been observed in *Neurospora, Arabidopsis* and human as well as *Drosophila* cell lines [3, 11-13]. This has provided convenient sequencing-based evidence of DNA break-induced antisense transcription.

While there is thus little doubt that a non-coding transcript initiates at the break (referred to as damage-induced long non-coding RNA or dilncRNA), we still do not have a comprehensive understanding of its biogenesis, in particular regarding whether differences exist between transcribed (i.e. within transcriptionally active genes) and non-transcribed breaks. *In vivo*, DNA breaks occur in a chromatin-context and the mechanisms of dilncRNA generation may differ depending on the local chromatin state, which determines the accessibility for RNA polymerases. For example, plants have even devoted the function of two polymerase II related, multi-subunit polymerases, RNA polymerase IV and IVb/V, to pervasive genome surveillance [14-17]. Their non-coding transcripts can activate a number of cellular responses to cope with transposon invasion, viral infection and also DNA breaks [13].

RNA polymerase I transcribes the rDNA and is largely confined to the nucleolus [18], whereas RNA polymerase III generates a series of non-coding transcripts. This polymerase also functions in certain cases to detect aberrant DNA: It transcribes AT-rich linear DNA that may be cytoplasmic [19, 20] or nuclear in the case of Herpesviruses [21-23]. The resulting pol-III transcripts then activate the cellular interferon response via RIG-I, an RNA helicase recognizing 5’-triphosphate-containing RNA in a double-stranded configuration [24]. Furthermore, RNA polymerase III can transcribe transposon-derived *Alu* elements and can direct new integration sites for the Ty1 transposon in budding yeast [25]. The transcriptional landscape of both, RNA pol II and pol III is thus complex and dynamic.

The notion that RNA polymerase II can initiate at a DNA break to generate a dilncRNA is supported by studies using RNA polymerase II specific inhibitors [26], chromatin-immunoprecipitation [27], single-molecule studies [10] and by the detection of dilncRNAs associated with RNA polymerase phosphorylated at tyrosine-1 within the CTD repeats in a metagene-analysis [28]. Recruitment of RNA polymerase II to the DNA end can involve the Mre11-Rad50-Nbs1 complex (MRN-complex) and transcription initiation at DNA breaks has indeed been reconstituted *in vitro* with linear DNA, purified RNA polymerase II and the MRN-complex [29]. The RNA polymerase II model has been challenged, however, by observations that claim selective recruitment of RNA polymerase III – also with the help of the MRN complex - to double-strand breaks in cultured human cells [30]. In *Drosophila*, the dilncRNA originating from a DNA break is converted into damage-induced siRNAs if the break occurs in actively transcribed genes [3]. The convergent transcripts form dsRNA, which is processed by the canonical RNAi machinery into Ago2-loaded siRNAs capable of silencing cognate transcripts [3, 31]. While their contribution to repair seems limited [31, 32], the siRNAs are much more stable than the original dilncRNA and thus can serve as a convenient proxy of dilncRNA transcription [3, 33]. Results from a genome-wide screen in *Drosophila* cells suggest that spliceosomes assembled on the normal transcript can stimulate the generation of corresponding damage-induced siRNAs. This was corroborated by the observation that DNA breaks upstream of a gene’s first intron or anywhere within intron-less genes produce few siRNAs upon damage [4].

An important question is thus whether the spliceosome acts upstream or downstream of the dilncRNA induction. In a downstream involvement, the spliceosome would serve as an RNA chaperone and promote the annealing of the coding (sense) and non-coding (antisense) transcripts, thereby boosting siRNA generation. An upstream action implies that the spliceosome stimulates the generation of dilncRNAs, i.e. recruitment of the polymerase to the break, and thereby increases the amount of dsRNA generated. We could now distinguish the two mechanisms by examining nascent transcription at DNA breaks and observed that a DSB downstream of introns leads to higher levels of antisense transcription, arguing that the spliceosome stimulates dilncRNA production. Furthermore, we propose that in *Drosophila* cells it is RNA polymerase II that transcribes the dilncRNA.

## Results and Discussion

The aim of our study was to measure the rate of antisense transcription at a transcribed DNA break for an intron-containing and an intronless gene. Furthermore, we wanted to determine which RNA polymerase is recruited for this purpose in *Drosophila*. Incorporation of labeled nucleotide analogs such as 4SU (4SU-Seq) or biotinylated dNTPs (PRO-Seq) allows to measure nascent transcriptomes with high sensitivity but cannot distinguish between RNA polymerases. While specific inhibitor treatments are available, they have the caveat that inhibition of RNA polymerase II will also abrogate transcription of the normal mRNA transcripts, which recruit the spliceosome and may thus participate in induction of antisense transcription at intron-containing genes. Yet, this is precisely what we wanted to test.

We therefore established a nascent RNA sequencing strategy based on polymerase-specific immunoprecipitation (nascent elongating transcript sequencing or NET-seq [34, 35]). In short, we lysed cultured *Drosophila* S2-cells harboring epitope-tags on RNA polymerase II or III (introduced via genome editing) and washed out cytoplasmic and soluble nuclear components. Then, a brief digestion with benzonase liberated chromatin-associated material (“input” in our figures), from which we could subsequently immunopurify tagged polymerases (“IP” in our figures). The short RNA stump protected by the polymerase during the benzonase treatment can directly enter our established small RNA sequencing library pipeline because benzonase products carry a 5’-monophosphate (see Fig S1 for an outline of our cell fractionation and NET-seq procedure). To verify our protocol, we sequenced both the input material for the IP (roughly speaking chromatin-associated RNA) and the polymerase-associated transcripts after immunoprecipitation.

### Validation of the NET-Seq procedure

We first examined the highly transcribed, protein-coding actin gene *act5C*. The profile of matching reads from the input material is dominated by the exonic portions of the gene, consistent with the notion that splicing can occur co-transcriptionally before release from the chromatin. Nonetheless, a certain level of intronic reads is already visible and demonstrates that the material also contains nascent transcripts. The nascent, RNA polymerase II associated reads sequenced after specific immunoprecipitation (IP) show a much stronger proportion of these intronic reads (Fig. 1 A, top panel). In comparison, the RNA polymerase III IP only showed non-specific background (distribution essentially unchanged - Fig. 1A, middle panel). Many genes show RNA polymerase II pausing shortly after transcript initiation. In *Drosophila*, this phenomenon was first comprehensively described in ChIP-Seq and PRO-seq experiments [36, 37]. Accordingly, promoter-proximal pausing is evident in the PRO-Seq trace for *act5C* as well as in our nuclear RNA sample (input) and particularly in the RNA-polymerase-II associated, nascent transcripts. When comparing our NET-Seq results for this highly abundant mRNA with published results of a nascent RNA labeling apporach (PRO-seq), it appears that our libraries still contain a moderate overrepresentation of exonic reads [38], presumably reflecting a higher background level in our NET-seq approach.

**Figure 1:**
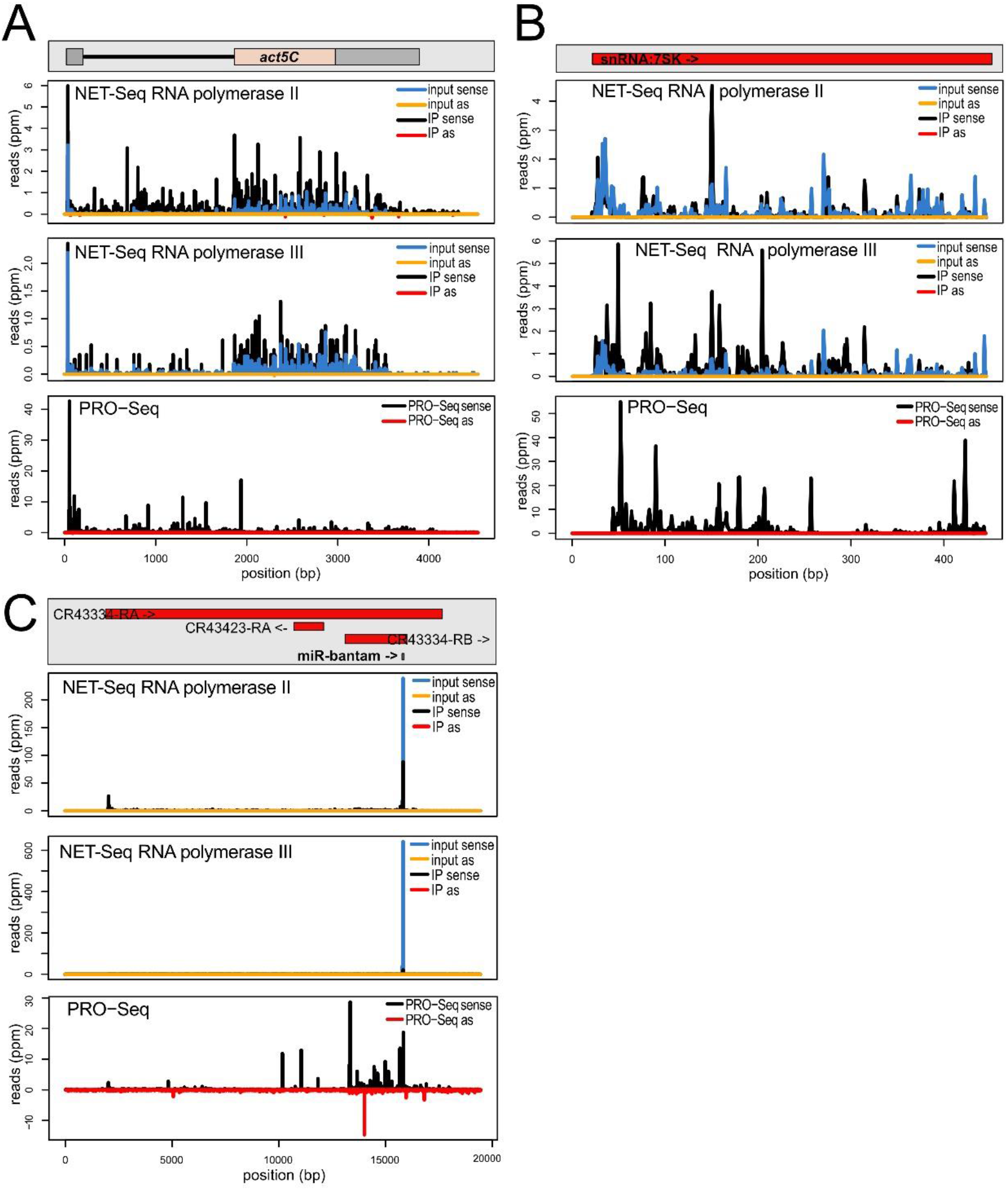
Characterization of the NET-Seq approach in *Drosophila* S2-cells A) NET-Seq reads for RNA polymerase II (top) and RNA polymerase III (middle) were mapped to the protein-coding gene *actin5C*; “input” refers to a chromatin-associated RNA fraction isolated prior to the polymerase-specific immunoprecipitation (IP). Reads from a published PRO-Seq experiment are shown at the bottom. B) NET-Seq reads for RNA polymerase II (top) and RNA polymerase III (middle) were mapped to the non-coding 7SK RNA gene, a known RNA polymerase III target; reads from a published PRO-Seq experiment are shown at the bottom. C) NET-Seq reads for RNA polymerase II (top) and RNA polymerase III (middle) were mapped to the *bantam* locus, an RNA polymerase II transcribed non-cding RNA; the mature *bantam* miRNA accumulates to high levels in the cytoplasm and is also an abundant contamination in our nuclear RNA preparations. Reads from a published PRO-Seq experiment are shown at the bottom.

For a global perspective, we also mapped reads onto precompiled transcript classes (Flybase genome release 6.19) and determined the recovery (ratio of IP versus input after normalization to total genome matching reads in each library) for RNA polymerase II and III. The CDS collection corresponds to the protein coding part of the transcriptome (start to stop) and the recovery was clearly greater in the pol-II IP than in the pol-III IP (Fig S2 A, pol-II IP n=6, pol-III IP n=4). The intronic part of the transcriptome also showed a preferential recovery with pol-II, but a certain number of introns also trended towards a high recovery in both, the pol-II and the pol-III IP (Fig S2 B). Manual inspection of an arbitrary subset usually indicated the presence of non-coding RNAs such as snRNAs or snoRNAs in these introns.

To verify successful IP for RNA polymerase III, we analyzed the read distribution along the non-coding 7SK RNA locus (Fig. 1B). While RNA polymerase II associated nascent transcripts did not show a particular enrichment of signal along the locus (top panel), the corresponding reads were enriched after IP of RNA polymerase III (middle panel). Note that the 7SK RNA can be associated with RNA polymerase II while the CTD is phosphorylated by pTEF-b; this associated 7SK RNA could thus have co-purified and augmented the 7SK-mapping read number. However, this does not appear to contribute substantially to the RNA polymerase II IP signal. As expected, the PRO-Seq procedure also captured transcription of the RNA polymerase III transcribed 7SK locus (bottom panel). When we mapped the reads onto the Flybase collection of tRNA sequences, we found a preferential recovery for at least a subset of the tRNAs in the RNA polymerase III IP (Fig S2 C). This is also visible when we mapped the reads onto the Flybase collection of “all transcripts”, which despite its name only comprises the protein-coding and lncRNAs. Essentially all of these are transcribed by RNA polymerase II but the *Ntl* locus is a notable exception (Fig S2 D). This transcript appears pol-III transcribed according to our analysis, overlaps with an intron-containing Tyr-GTA tRNA gene and direct visualization of the mapping traces revealed that the read-counts mapped to the *Ntl* locus almost exclusively localize to the tRNA portion (Fig S2 E).

Our Net-Seq libraries are contaminated by abundant cytoplasmic non-coding RNAs. This is illustrated with the help of the *bantam* locus (Fig. 1C). The 23 nt small RNA is one of the most abundant miRNAs in S2-cells and it is nucleolytically processed from a much larger primary transcript by Drosha and Dicer-1. The mature miRNA is cytoplasmic, yet our nuclear RNA fraction still contained a substantial amount of bantam reads (top and middle panel, input). While the IP procedure decreased this contamination, it did not remove the bantam reads completely (top and middle panel, IP). However, in the case of RNA polymerase II the nascent RNA reads indicate that larger precursor ncRNAs are transcribed (top panel, IP). This is consistent with the PRO-Seq reads from the locus (bottom panel). The three example loci for Fig. 1 were chosen because the published PRO-Seq reads can be represented at roughly comparable ppm-scales, hence their transcriptional output should be, as a first approximation, of comparable magnitude. Our own Net-Seq data for *act5C* and *7SK* can indeed also be displayed with comparable scales, but the *bantam* locus required different scaling due to the cytoplasmic contamination. We also observed a substantial amount of mature ribosomal RNA reads in our libraries both, before and after IP (23%-72% of total genome-matching reads, with no obvious enrichment of unprocessed precursor transcripts). For these RNAs, no interpretation of our sequencing data should be attempted. This also limits conclusions about highly abundant RNAs transcribed by RNA polymerase III such as 5S rRNA. For most other transcripts, we conclude that our nascent RNA sequencing data successfully captures polymerase-specific profiles. Since our question focuses on the induced antisense transcription at DNA breaks, an RNA species that is neither cytoplasmic nor highly abundant, we conclude that the NET-Seq libraries are suitable for our analysis.

### A DSB downstream of introns shows higher dilncRNA transcription activity

We generated sequencing libraries after employing our established *cas9*/CRISPR system to cleave in the intron-containing gene *CG15098* and, separately, in the intronless gene *tctp* [4]. As before, the DNA breaks had been induced by transfection of a corresponding sgRNA expression cassette into cells that stably express the Cas9 protein. The majority of the cells were harvested and processed for NETseq libraries 2 or 3 days after transfection. The remaining cells were processed for a T7 endonuclease assay, demonstrating that the targeted loci were indeed cleaved with comparable efficiency (see also Fig S1). In our experiments, libraries from the *tctp*-cut provide the “uncut” control for the *CG15098* locus and *vice-versa*. This comparison ensures that any effects not specific to the cut locus or due to Cas9 activation *per se* will be accounted for.

We mapped the NET-seq libraries onto the respective loci and calculated the number of sense and antisense-matching reads. Figure 2 shows traces for one NET-Seq replicate mapped to *CG15098* (left side) and *tctp* (right side). For *CG15098*, IP of RNA polymerase II associated, nascent transcripts led to an enrichment of antisense reads relative to input (Fig. 2A). In contrast, the antisense reads did not increase for the cut *tctp* locus, consistent with the low amounts of siRNAs generated upon cleavage of this locus [4]. There was no indication for a prominent signal in the RNA polymerase III NET-seq libraries of either locus (Fig. 2B).

**Figure 2:**
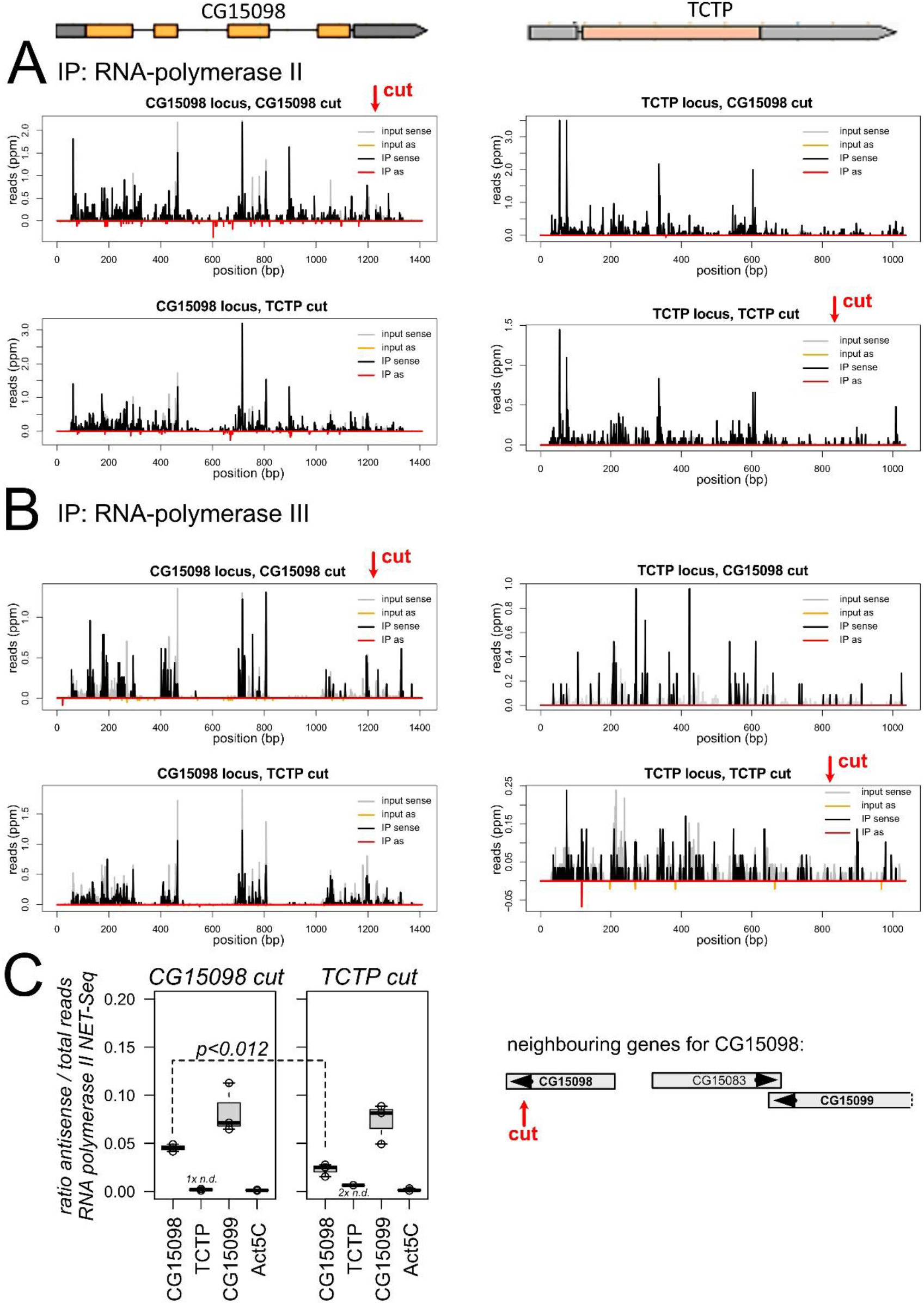
Net-Seq analysis of dilncRNA transcription A) Sample traces for one replicate showing the NET-seq reads of RNA polymerase II mapped to *CG15098* (left) and *tctp* (right). In the top row the *CG15098* locus was cleaved, while *tctp* was cleaved in the bottom row. B) Same as A but showing the NET-seq reads of RNA polymerase III. C) Quantitative analysis of the antisense reads relative to all reads mapped to the respective locus revealed a significant increase for *CG15098* in the cleaved state (left, t-test unequal variance, n=3). A cartoon shows the genes in the vicinity of *CG15098*, the closest neighbor in the same orientation is *CG15099*. Note that this gene is convergent with CG15083 and thus intrinsically has a higher proportion of antisense transcripts that map to the overlapping region.

To obtain a quantitative view of the replicate data, we normalized the number of antisense reads to the total transcriptional activity of the locus in each library [i.e. antisense /(sense + antisense)] (Fig. 2C). There was a significant increase of antisense reads for cut vs. uncut *CG15098* (*p*=0.012, t-test unpaired, unequal variance, n=3) while no significant differences were observed for the neighboring *CG15099* (*p*=0.640, n=3) or *act5C*, which resides on a different chromosome (*p*=0,644, n=3). We also normalized the antisense reads to the total number of genome-matching reads in each library (Fig EV 3). In each of the three replicate experiments, the amount of *CG15098* antisense-matching nascent, RNA polymerase II associated reads was higher in the cut state than in the uncut state (*p*=0.034, paired t-test, n=3). This was not the case for *CG15099* gene (*p*=0.273, n=3) or the *act5C* gene (*p*=0.675, n=3); there were too few *tctp* antisense matching reads for an analogous comparison. Finally, our input material also showed a consistently higher amount of antisense-matching reads for *CG15098* in the cut state in each replicate (*p*=0.072, paired t-test, n=3). In agreement with the visual inspection (Fig. 2B), the read quantification did not provide any indication that RNA polymerase III is contributing to antisense transcription (Fig S3, bottom row).

We conclude that induction of a DNA double-strand break in the intron-containing *CG15098* gene stimulates antisense transcription by RNA polymerase II. For the intronless *tctp*-gene, we detected none or only few antisense reads and statistical analysis is not appropriate. Our observations are thus consistent with the notion that a lower antisense transcription activity for the intronless gene (this study) correlates with fewer DNA-damage induced siRNAs [4]. It therefore appears that the role of the spliceosome is to stimulate dilncRNA transcription, rather than to promote annealing of the sense and antisense RNA strands. Since the overwhelming majority of fly genes contains at least one intron, many spontaneously occurring DSB’s can be affected by this phenomenon.

Our finding also has important mechanistic implications since it could be the very same polymerase that synthesizes both sense and antisense transcript. In this most rudimentary form of “recruitment”, stalling of the splicing reaction could e.g. contribute to post-transcriptional modifications on RNA polymerase II that promote direct re-initiation upon a run-off at the break – a “U-turn” movement, essentially. However, it is currently unclear whether a run-off will occur at a DSB *in vivo* or whether the polymerase stalls when it encounters the break. As long as the transcript is not cleaved and removed, this creates an R-loop behind the polymerase with concomitant exposure of the non-template strand. This stretch of single-stranded DNA could also serve as a landing site for another RNA polymerase complex and transcription thus initiates in the antisense orientation. In this case, the role of the stalled spliceosome could be to prevent transcript termination and release, thus extending the lifetime of the R-loop that may contribute to DNA damage signaling. Alternatively or in addition to this modulation of the DNA/RNA duplex structures, signaling events that include or emanate from spliceosome components [39] could foster polymerase recruitment to the nearby single-stranded DNA.

### No evidence for participation of RNA polymerase III in the biogenesis of damage-induced siRNAs

The recent description of RNA polymerase III recruitment to DNA breaks in human cell lines [30] clearly differs from our observation of a predominant – if not exclusive - role of RNA polymerase II in dilncRNA generation (Fig. 2). It is certainly conceivable that mechanistic differences exist between humans and flies (as is the case for the subsequent processing into siRNAs, see [32]), but we wanted to confirm our observation with independent approaches. We thus turned to our established dual luciferase reporter system, which relies on the silencing activity of damage-induced siRNAs generated from a co-transfected, linearized plasmid (Fig. 3A, right side). With this assay, we had previously screened and detected a role for the MRN-complex in promoting siRNA generation, presumably by preparing the DNA end for RNA polymerases that initiate transcription at the break [4]. The inhibitor Mirin can block the access of Mre11 to dsDNA ends and thus all nucleolytic activities, while its derivative PFM-01 selectively blocks DNA access to the endonuclease active site [40]. Addition of Mirin (25 µM final concentration) clearly reduced the amount of damage-induced siRNAs generated (*p*=0.05, t-test, unequal variance, n=3), while PFM-01 (25 µM) had essentially no effect (Fig. 3A). This supports the notion that the initial unwinding of the double-stranded DNA by Mre-11 is important for dilncRNA generation, rather than endonucleolytic cleavage and resection that exposes single-stranded DNA with a 3’-end [29].

**Figure 3:**
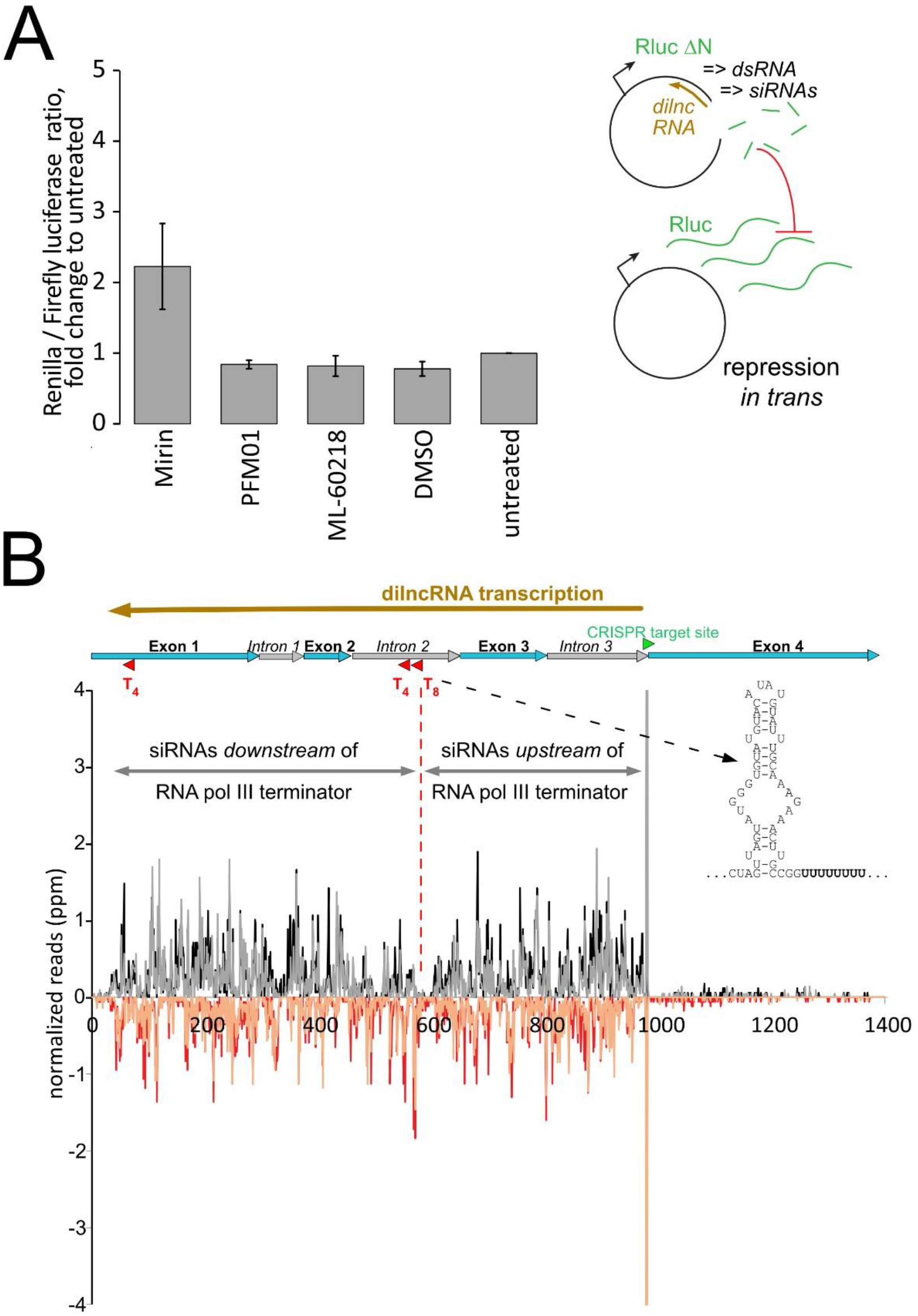
No alternative evidence for participation of RNA polymerase III in dilncRNA transcription A) Luciferase-encoding plasmid based assay for the detection of damage-induced siRNAs; a linearized plasmid with a truncated *Renilla* luciferase gene serves as donor for dilncRNA transcription, dsRNA formation and processing into siRNAs. These in turn repress a co-transfected full-length *Renilla* luciferase vector. Inhibition of the MRN-complex with the inhibitor Mirin, but not PFM-01, reduced the amount of damage-induced siRNAs. Inhibition of RNA polymerase III with ML-60218, however, did not lead to any change of siRNA yield compared with the solvent control (DMSO). Three biological replicates of the assay were performed. B) A stretch of 8 adenosines in the second intron of *CG15098* will lead to a corresponding sequence of 8 thymines in the dilncRNA transcript. This is preceded by a potential secondary structure element (shown on the right in 5’->3’ direction of an antisense transcript) and should lead to termination of RNA polymerase III transcription. Hence, a lower density of damage-induced siRNAs should be observed beyond this point if RNA polymerase III transcribes the dilncRNA. This was, however, not the case. (sequencing data previously published in [4]).

**Figure 4:**
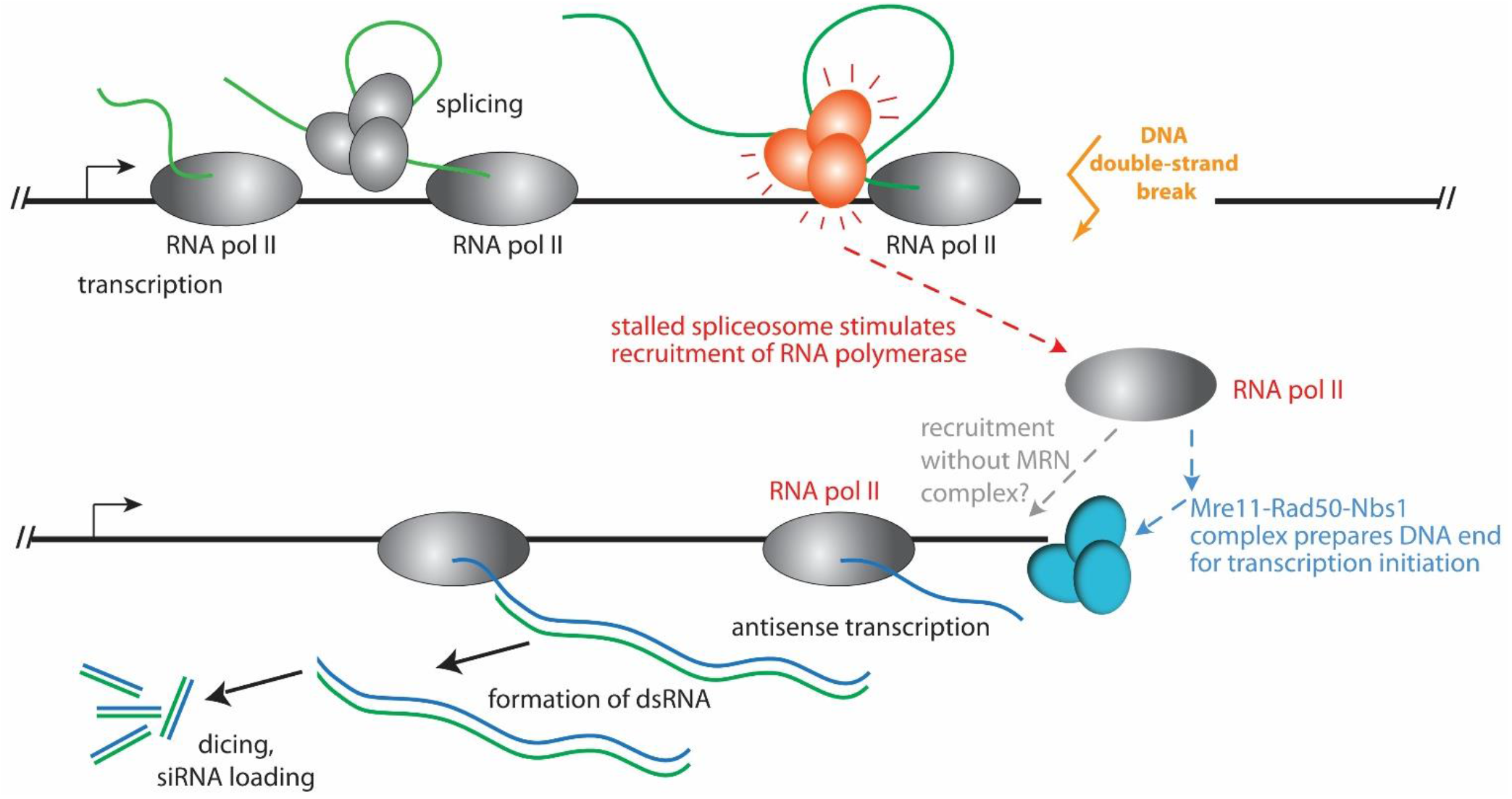
Model for the recruitment of RNA polymerase II to a DNA double-strand break in *Drosophila* Our new results demonstrate that the role of the spliceosome is to recruit RNA polymerase II for antisense transcription (red arrow). This is a key progress because we can now rule out a mere RNA-chaperone-like activity of the spliceosome. Rather, local transcription and splicing are important mediators of dilncRNA biogenesis. As previously described for mammalian cells, transcript initiation at the break is aided by the Mre11-Rad50-Nbs1 complex in *Drosophila* as well. Future experiments can address whether the spliceosome-mediated recruitment requires MRN, or whether the two pathways are independent possibilities for recruiting RNA polymerase II to the DNA break. While our NET-seq approach, inhibitor treatments and sequence analysis of the *CG15098* model locus did not provide any evidence for RNA polymerase III mediated dilncRNA transcription, we cannot rule out the possibility that RNA polymerase III is recruited to the break without engaging in processive transcription.

Importantly, addition of the selective RNA polymerase III inhibitor ML-60218 at a concentration of 10 µM - the highest concentration that still produced acceptable levels of luciferase readings - did not lead to a de-repression of *Renilla* luciferase (Fig. 3A). This is consistent with our genome-wide RNAi screen where no RNA polymerase III subunit scored as a confirmed hit [4] and with undetectable dilncRNA transcription in our RNA pol-III NET-seq libraries.

We had previously determined that the damage-induced siRNA response starts in close proximity to the break and extends all the way until the transcription start site [3, 4, 33]. The corresponding dilncRNA transcripts thus arise over a stretch of more than 1 kb (e.g. 4.5 kb in the case of CG18273, see supplementary Figures in [4]). This would be unusually long for an RNA polymerase III transcript and random pol-III termination sequences might occur along the way. Indeed, inspection of the *CG15098* locus revealed a serendipitous stretch of eight Adenosines in the second intron. For an RNA polymerase acting in antisense orientation, this corresponds to a T_8_-sequence preceded by a potential secondary structure element (see Fig. 3B), which should terminate most RNA polymerase III transcription complexes [41]. However, the siRNA read density we observed was similar before and after this pol-III termination site (Fig. 3B). We do note that there is a paucity of siRNA reads in a ∼ 20 nt window surrounding the A_8_/T_8_ sequence; most likely this is for technical reasons given the short, homopolymeric sequence stretch (e.g. Illumina-sequencing or PCR polymerase drop-off). While it is thus unlikely that RNA polymerase III functionally contributes to dilncRNA transcription in *Drosophila*, our observations cannot exclude that RNA polymerase III is recruited to sites of DNA damage without subsequently engaging in processive transcription of the dilncRNA.

Previously, several publications have provided independent evidence of RNA polymerase II as an enzyme capable of transcribing the dilncRNA. This includes biochemical reconstitutions [29], *in vitro* analysis with inhibitors [26], ChIP with qPCR [26] and metagene analysis after ChIP-Seq [28]. A single-molecule study is also suggestive of RNA polymerase II according to the reported speed [10], but the MS2 stem-loop employed as a reporter can in principle also be transcribed by RNA polymerase III [42]. While not all of the published experiments can exclude a concomitant function of more than one RNA polymerase - i.e. RNA polymerase II (or IV in plants) *and* RNA polymerase III - in dilncRNA generation, the recent description of RNA polymerase III as the exclusive source of dilncRNA in cultured human cell lines is surprising [30]. We now provide a direct observation of polymerase-associated, nascent dilncRNA transcripts only in RNA polymerase II NET-seq. Certainly, differences between organisms may exist: If the primary purpose is to generate a transcript, then the polymerase type could easily be swapped during the course of evolution. In plants, for example, genetic analysis has pinpointed a function of the plant-specific RNA polymerase IV in dilncRNA transcription [13]. The situation is further complicated by the discovery that repair of transcribed genes by homologous recombination is also fostered upon the establishment of mixed DNA/RNA displacement loops involving the normal transcript that runs sense towards the break [43]. A parallel comparison of the diverse experimental systems might help to distinguish between technical and true biological differences; the latter will prove invaluable to further our understanding of the molecular mechanisms that lead to dilncRNA transcription.

## Materials and Methods

### NET-Seq procedure

#### Cell culture

*Drosophila* S2-cells with stable expression of cas9 protein (clone 5-3) were cultured and transfected as previously described [44]. We further modified this cell line by introducing a twin V5-tag at the C-terminus of the largest subunit of RNA polymerase II (PolR2A, CG1554) and III (PolR3A, CG17209), followed by clonal selection as described [45]. For the NET-seq experiments, we transfected a 30 ml culture of cells expressing tagged RNA polymerase with guideRNA vectors targeting *CG15098* or *tctp*. The sgRNA expression cassettes were first generated by PCR, then blunt-end cloned into pJet1.2 to yield pRB59 (*CG15098*) and pRB60 (*tctp*). The target sites were 5’-TCCAGTGTAGCTTCCCGTT-3’ for *CG15098* and 5’-ATATCTAATTTCTTTTTAC-3’ for *tctp* as described [4].

#### Cell lysis

48 or 36 hours after transfection, the cells were harvested (density 4-5 × 10^6^ cells/ml), resuspended in 500 µl of lysis buffer (10 mM HEPES/KOH PH7.5, 1.5 mM MgCl_2_, 1 mM DTT, 10 mM EDTA, 10% glycerol and 1% Tergitol-type NP40 (Sigma NP40S) supplemented with proteinase inhibitors (Roche complete without EDTA)) and incubated for 10 minutes on ice. Then nuclei were pelleted by centrifugation at 5000xg for 5 minutes and the supernatant (mostly cytosol) was discarded. The pellet was resuspended in lysis buffer without EDTA but containing 1 M urea, incubated for 5 minutes on ice and again pelleted at 5000xg for 5 minutes. The urea washing step was carried out twice in total, then the nuclei were resuspended in 110 µl of lysis buffer without EDTA and without urea. To digest the chromatin, 250 U of benzonase (Merck Millipore E1014, 90% purity grade) were added and the resuspended nuclei were incubated at 37°C for 3 minutes in a heating block. The digestion was stopped by adding EDTA and NaCl to a concentration of 10 mM and 500 mM, respectively. The insoluble fraction was pelleted by centrifugation at 16000xg for 5 minutes and the supernatant was used as input material for the immunoprecipitation.

#### Immunoprecipitation

20 µl of magnetic beads (Dynabeads protein G, Invitrogen 10004D) were washed 3 times with 200 µl of IP buffer (25 mM HEPES/KOH pH 7.5, 150 mM NaCl, 12.5 mM MgCL2, 1 mM DTT, 1% Tergitol-Type NP40, 0.1% Empigen (Sigma 30326) supplemented with Roche complete proteinase inhibitors without EDTA), then 1 µl of V5 antibody was coupled by rotation at 4 °C over night. On the following day, the beads were washed 3x with 300 µl of IP buffer, then the input material was added and incubated with agitation for 60 minutes at 4°C. After separation of the unbound supernatant, the beads were washed 5x with 200 µl of IP-buffer. The immunopurified RNA polymerase complexes were the digested with proteinase K to liberate the associated nucleic acids and RNA was prepared by TRIZOL extraction and precipitation.

#### Library generation and data analysis

RNA fragments with a size of 20-28 nt were PAGE-purified to select for the fragments that were protected from benzonase digestion by the polymerase. Since benzonase products harbor 5’-phosphorylated ends, the RNA fragments were processed for library generation as described [46] without further treatment. The libraries were sequenced in-house on an Illumina HiSeq1500 instrument and the reads were processed with custom PERL and BASH scripts for mapping with Bowtie [47] to the indicated references. During mapping, no mismatches were tolerated and each hit was reported only once. If multiple, perfectly matching sequences exist in the reference, the Bowtie algorithm will assign the read randomly. After mapping, the results were further processed with BEDtools [48] and custom R!-scripts or the IGV genome browser [49] for data visualization.

### Luciferase assay

The luciferase assay for the detection of DNA-break induced siRNAs has been previously described [4]. Briefly, 25 ng of pRB2 (firefly-luciferase, circular), 10 ng of pRB1 (Renilla-luciferase, circular) and 40 ng of pRB4 (truncated Renilla luciferase, linearized with EcoRI) were transfected per well of a 96-well plate using Fugene-HD (Promgea). Inhibitors were added 2 hr prior to transfection in a volume of 1 µl DMSO (volume identical for all compounds and controls). The luciferase assay was performed 96 hrs after transfection using the Dual-Glo Luciferase assay system (Promega E2920) in a Tecan M-1000 plate reader. Data analysis was carried out using Microsoft Excel.

## Data availability

The sequencing reads from this study are available at the European Nucleotide Archive with the accession number PRJEB12939. Custom PERL, BASH and R! scripts have been deposited on Github: https://github.com/Foerstemann/small_RNA_seq_analysis.git

## Acknowledgements

We are grateful for the sequencing support by the laboratory of functional genome analysis (LaFuGa) at the Gene Center Munich. This study has been supported by DFG grant FO-360/9-1 to KF.

## Author contributions

RB, PD and IS performed experiments, VN provided essential reagents, KF, RB and IS designed experiments, all authors wrote and edited the manuscript. KF conceived the project and provided funding.

## Supplementary Figure legends

**Figure S1:**
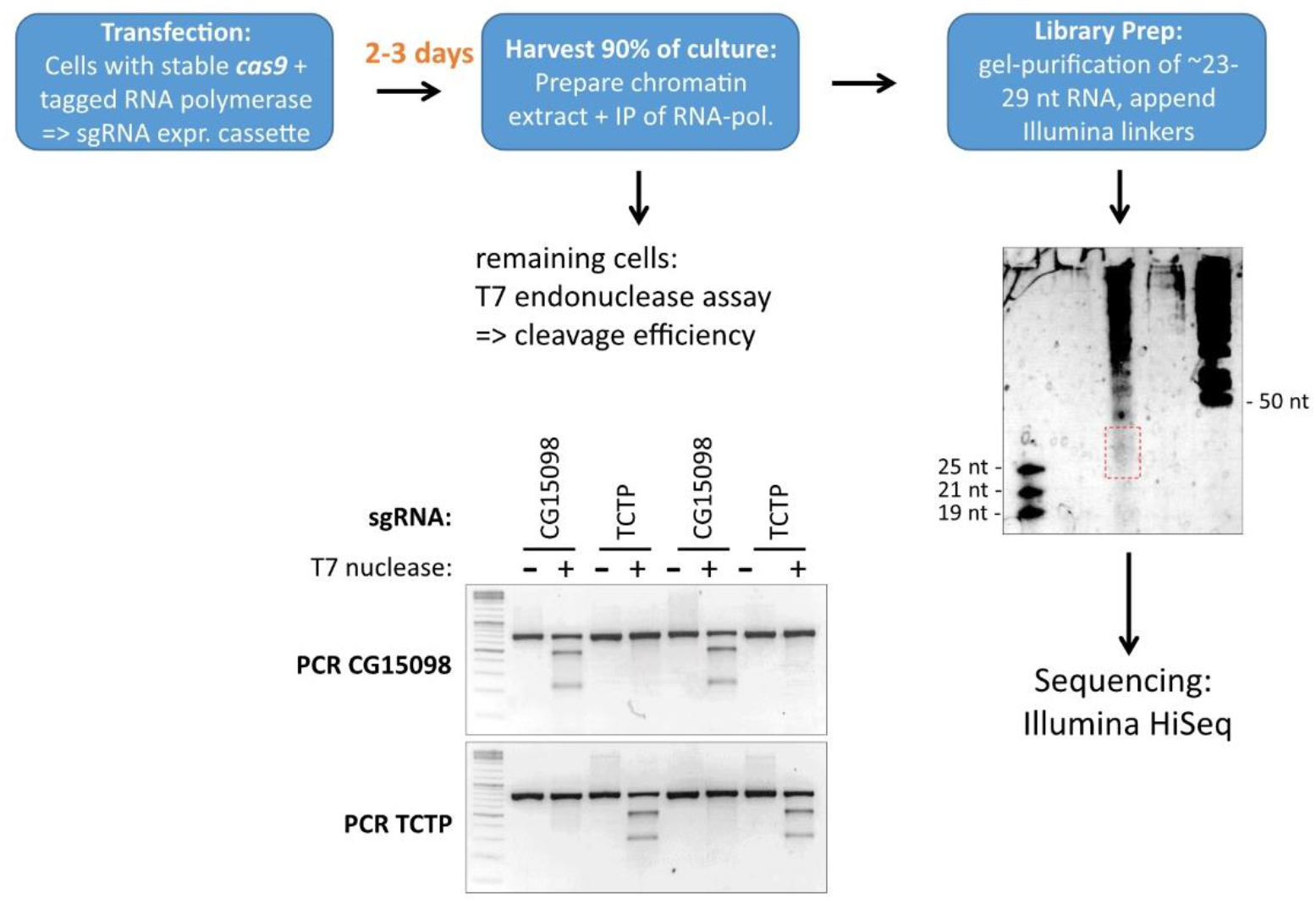
Outline of the NET-seq procedure and sample gel of quality control for cleavage efficiency

**Figure S2:**
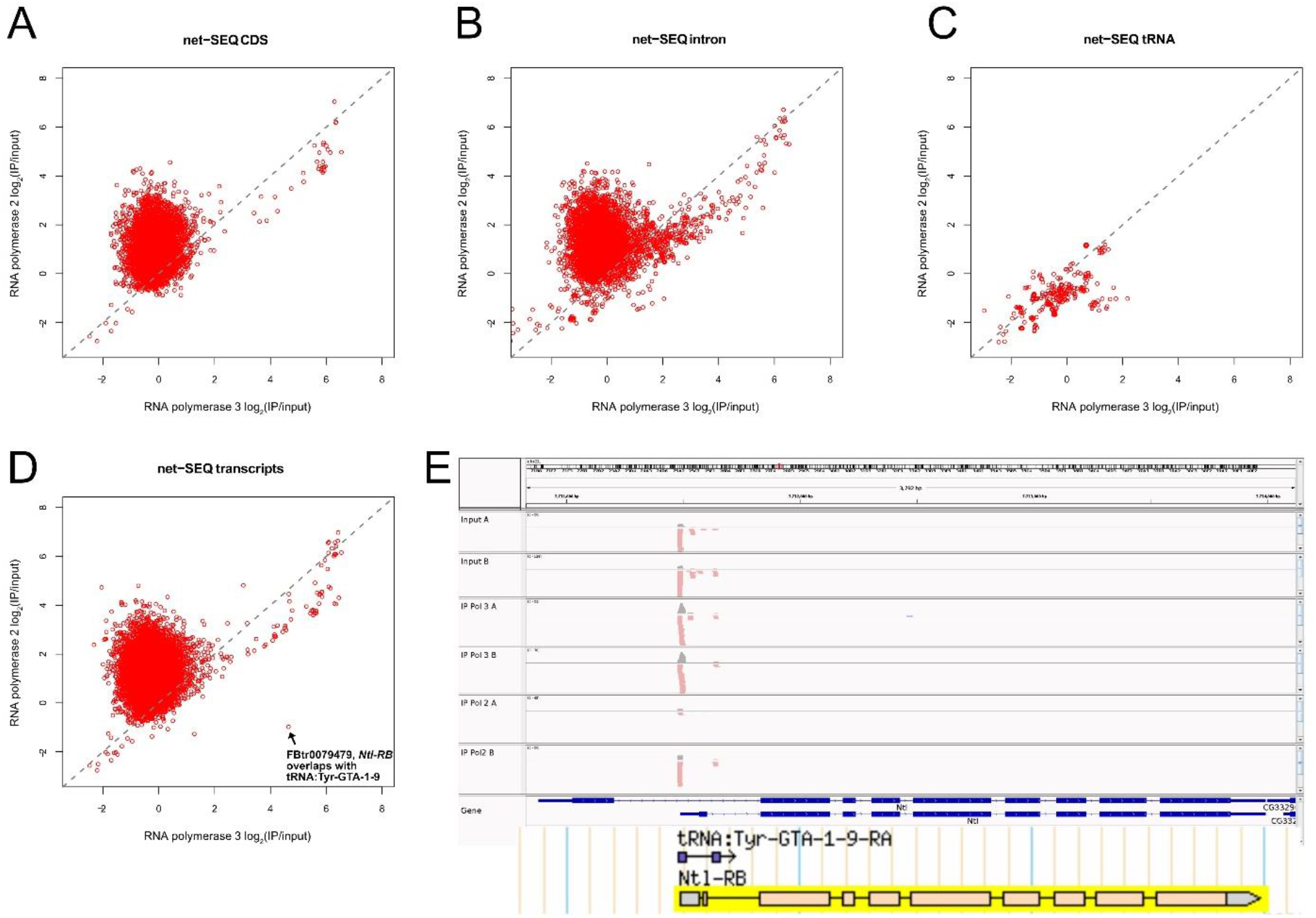
Genome-wide enrichment analysis for RNA polymerase II and III A) Mapping all reads to the precompiled protein-coding sequences (= ATG-to-stop) reveals selective enrichment upon IP of RNA polymerase II B) Mapping all reads to the precompiled intron sequences reveals enrichment upon IP of RNA polymerase II in many cases, but a number of introns appear to be at least in part transcribed by RNA polymerase III. Manual inspection of a subset indicates that these often harbor highly abundant non-coding RNA genes (snRNAs, snoRNAs etc.); it is unclear whether this shows *bona fide* pol III transcription or contamination by abundant RNA species. C) At least a subset of tRNA genes is clearly enriched upon IP of RNA polymerase III. D) Mapping all reads to the precompiled “all transcripts” collection (= protein-coding and non-coding RNA polymerase II transcripts) reveals selective enrichment upon IP of RNA polymerase II with the notable exception of the *Ntl* locus. E) The *Ntl* locus harbors a Tyr-GTA tRNA gene in the first intron, the mapping traces demonstrate that the assigned reads only come from the tRNA gene and that they are clearly enriched in RNA polymerase III NET-seq libraries (tracks scaled according to total genome matching reads in each library). Plase note that there are 6 Tyr-GTA tRNA genes in the fly genome with identical mature sequence and only the gene within the *Ntl* locus contains an intron. Because of the identical sequence, transcripts arising from any of the 6 tRNA loci will be mapped to each of the 6 genes. We thus cannot conclude that the *Ntl* locus, which fortuitously harbors a tRNA gene, is pol-III transcribed, but only that at least one of the Tyr-GTA tRNA loci is pol-III transcribed.

**Figure S3:**
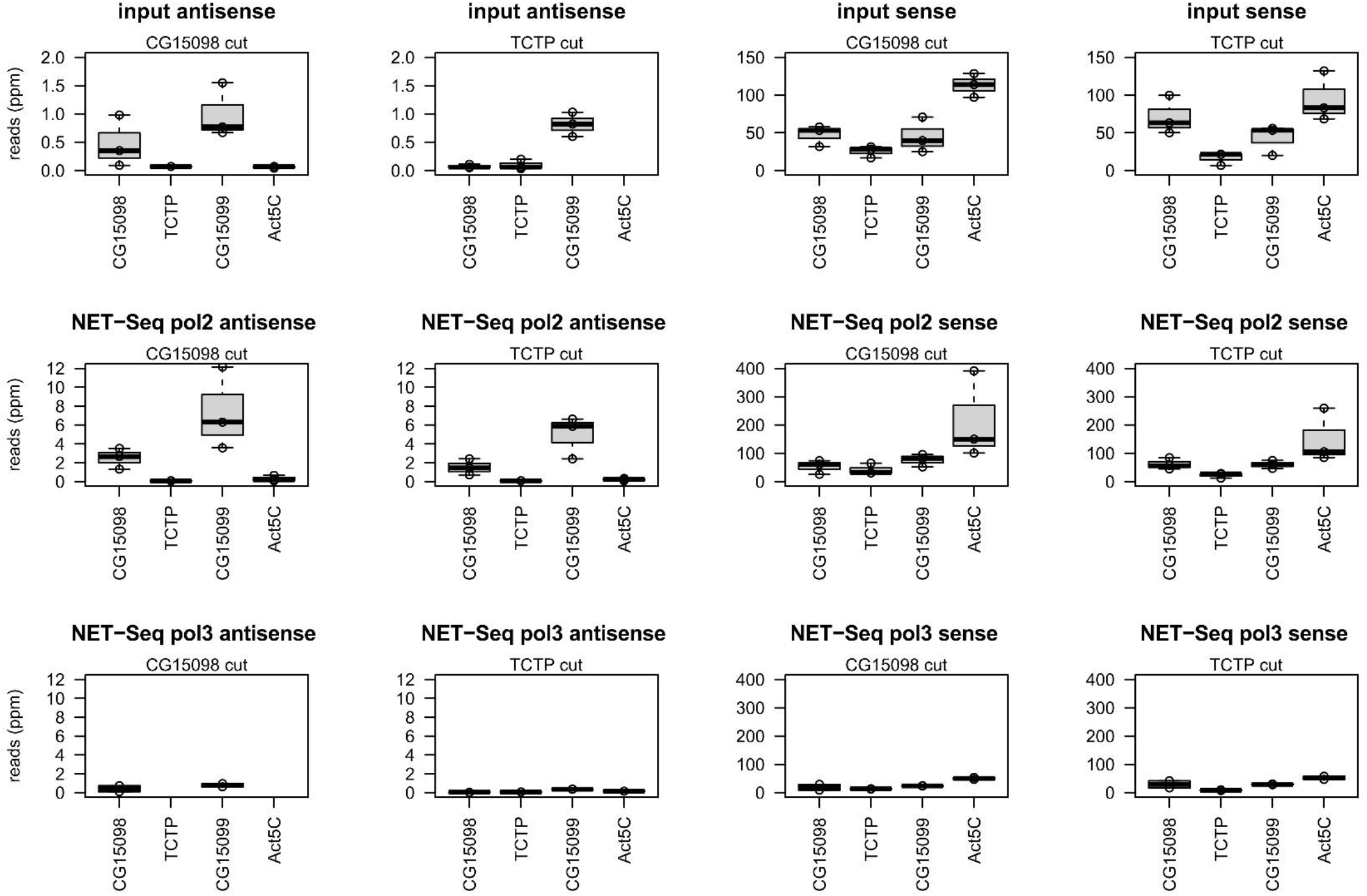
Quantification of NET-seq and input material for a set of genes (*CG15098*, TCTP, *CG15099* and Act5C)

## References

1. van Rij, R.P., M.C. Saleh, B. Berry, C. Foo, A. Houk, C. Antoniewski, et al., The RNA silencing endonuclease Argonaute 2 mediates specific antiviral immunity in Drosophila melanogaster. Genes & development, 2006. 20(21): p. 2985–95.

2. Wood, J.G., B.C. Jones, N. Jiang, C. Chang, S. Hosier, P. Wickremesinghe, et al., Chromatin-modifying genetic interventions suppress age-associated transposable element activation and extend life span in Drosophila. Proc Natl Acad Sci U S A, 2016. 113(40): p. 11277–11282.

3. Michalik, K.M., R. Bottcher, and K. Forstemann, A small RNA response at DNA ends in Drosophila. Nucleic acids research, 2012. 40(19): p. 9596–603.

4. Merk, K., M. Breinig, R. Bottcher, S. Krebs, H. Blum, M. Boutros, et al., Splicing stimulates siRNA formation at Drosophila DNA double-strand breaks. PLoS Genet, 2017. 13(6): p. e1006861.

5. Dumesic, P.A., P. Natarajan, C. Chen, I.A. Drinnenberg, B.J. Schiller, J. Thompson, et al., Stalled spliceosomes are a signal for RNAi-mediated genome defense. Cell, 2013. 152(5): p. 957–68.

6. Ciccia, A. and S.J. Elledge, The DNA damage response: making it safe to play with knives. Mol Cell, 2010. 40(2): p. 179–204.

7. Caron, P., J. van der Linden, and H. van Attikum, Bon voyage: A transcriptional journey around DNA breaks. DNA Repair (Amst), 2019. 82: p. 102686.

8. Ohle, C., R. Tesorero, G. Schermann, N. Dobrev, I. Sinning, and T. Fischer, Transient RNA-DNA Hybrids Are Required for Efficient Double-Strand Break Repair. Cell, 2016. 167(4): p. 1001–1013 e7.

9. Iannelli, F., A. Galbiati, I. Capozzo, Q. Nguyen, B. Magnuson, F. Michelini, et al., A damaged genome’s transcriptional landscape through multilayered expression profiling around in situ-mapped DNA double-strand breaks. Nat Commun, 2017. 8: p. 15656.

10. Vitor, A.C., S.C. Sridhara, J.C. Sabino, A.I. Afonso, A.R. Grosso, R.M. Martin, et al., Single-molecule imaging of transcription at damaged chromatin. Sci Adv, 2019. 5(1): p. eaau1249.

11. Lee, H.-C., S.-S. Chang, S. Choudhary, A.P. Aalto, M. Maiti, D.H. Bamford, et al., qiRNA is a new type of small interfering RNA induced by DNA damage. Nature, 2009. 459(7244): p. 274–277.

12. Francia, S., F. Michelini, A. Saxena, D. Tang, M. de Hoon, V. Anelli, et al., Site-specific DICER and DROSHA RNA products control the DNA-damage response. Nature, 2012. 488(7410): p. 231–5.

13. Wei, W., Z. Ba, M. Gao, Y. Wu, Y. Ma, S. Amiard, et al., A Role for Small RNAs in DNA Double-Strand Break Repair. Cell, 2012. 149(1): p. 101–12.

14. Herr, A.J., M.B. Jensen, T. Dalmay, and D.C. Baulcombe, RNA polymerase IV directs silencing of endogenous DNA. Science, 2005. 308(5718): p. 118–20.

15. Wierzbicki, A.T., J.R. Haag, and C.S. Pikaard, Noncoding transcription by RNA polymerase Pol IVb/Pol V mediates transcriptional silencing of overlapping and adjacent genes. Cell, 2008. 135(4): p. 635–48.

16. Pontier, D., G. Yahubyan, D. Vega, A. Bulski, J. Saez-Vasquez, M.A. Hakimi, et al., Reinforcement of silencing at transposons and highly repeated sequences requires the concerted action of two distinct RNA polymerases IV in Arabidopsis. Genes Dev, 2005. 19(17): p. 2030–40.

17. Tsuzuki, M., S. Sethuraman, A.N. Coke, M.H. Rothi, A.P. Boyle, and A.T. Wierzbicki, Broad noncoding transcription suggests genome surveillance by RNA polymerase V. Proc Natl Acad Sci U S A, 2020. 117(48): p. 30799–30804.

18. Scheer, U. and K.M. Rose, Localization of RNA polymerase I in interphase cells and mitotic chromosomes by light and electron microscopic immunocytochemistry. Proc Natl Acad Sci U S A, 1984. 81(5): p. 1431–5.

19. Ablasser, A., F. Bauernfeind, G. Hartmann, E. Latz, K.A. Fitzgerald, and V. Hornung, RIG-I-dependent sensing of poly(dA:dT) through the induction of an RNA polymerase III-transcribed RNA intermediate. Nat Immunol, 2009. 10(10): p. 1065–72.

20. Chiu, Y.H., J.B. Macmillan, and Z.J. Chen, RNA polymerase III detects cytosolic DNA and induces type I interferons through the RIG-I pathway. Cell, 2009. 138(3): p. 576–91.

21. Ogunjimi, B., S.Y. Zhang, K.B. Sorensen, K.A. Skipper, M. Carter-Timofte, G. Kerner, et al., Inborn errors in RNA polymerase III underlie severe varicella zoster virus infections. J Clin Invest, 2017. 127(9): p. 3543–3556.

22. Yu, H.R., H.C. Huang, H.C. Kuo, J.M. Sheen, C.Y. Ou, T.Y. Hsu, et al., IFN-alpha production by human mononuclear cells infected with varicella-zoster virus through TLR9-dependent and -independent pathways. Cell Mol Immunol, 2011. 8(2): p. 181–8.

23. Ramanathan, A., M. Weintraub, N. Orlovetskie, R. Serruya, D. Mani, O. Marcu, et al., A mutation in POLR3E impairs antiviral immune response and RNA polymerase III. Proc Natl Acad Sci U S A, 2020. 117(36): p. 22113–22121.

24. Hornung, V., J. Ellegast, S. Kim, K. Brzozka, A. Jung, H. Kato, et al., 5’-Triphosphate RNA is the ligand for RIG-I. Science, 2006. 314(5801): p. 994–7.

25. Asif-Laidin, A., C. Conesa, A. Bonnet, C. Grison, I. Adhya, R. Menouni, et al., A small targeting domain in Ty1 integrase is sufficient to direct retrotransposon integration upstream of tRNA genes. EMBO J, 2020. 39(17): p. e104337.

26. Michelini, F., S. Pitchiaya, V. Vitelli, S. Sharma, U. Gioia, F. Pessina, et al., Damage-induced lncRNAs control the DNA damage response through interaction with DDRNAs at individual double-strand breaks. Nat Cell Biol, 2017. 19(12): p. 1400–1411.

27. D’Alessandro, G., D.R. Whelan, S.M. Howard, V. Vitelli, X. Renaudin, M. Adamowicz, et al., BRCA2 controls DNA:RNA hybrid level at DSBs by mediating RNase H2 recruitment. Nat Commun, 2018. 9(1): p. 5376.

28. Burger, K., M. Schlackow, and M. Gullerova, Tyrosine kinase c-Abl couples RNA polymerase II transcription to DNA double-strand breaks. Nucleic Acids Res, 2019. 47(7): p. 3467–3484.

29. Sharma, S., R. Anand, X. Zhang, S. Francia, F. Michelini, A. Galbiati, et al., MRE11-RAD50-NBS1 Complex Is Sufficient to Promote Transcription by RNA Polymerase II at Double-Strand Breaks by Melting DNA Ends. Cell Rep, 2021. 34(1): p. 108565.

30. Liu, S., Y. Hua, J. Wang, L. Li, J. Yuan, B. Zhang, et al., RNA polymerase III is required for the repair of DNA double-strand breaks by homologous recombination. Cell, 2021. 184(5): p. 1314–1329 e10.

31. Schmidts, I., R. Bottcher, M. Mirkovic-Hosle, and K. Forstemann, Homology directed repair is unaffected by the absence of siRNAs in Drosophila melanogaster. Nucleic acids research, 2016. 44(17): p. 8261–71.

32. Bonath, F., J. Domingo-Prim, M. Tarbier, M.R. Friedlander, and N. Visa, Next-generation sequencing reveals two populations of damage-induced small RNAs at endogenous DNA double-strand breaks. Nucleic Acids Res, 2018. 46(22): p. 11869–11882.

33. Kunzelmann, S. and K. Forstemann, Reversible perturbations of gene regulation after genome editing in Drosophila cells. PLoS One, 2017. 12(6): p. e0180135.

34. Churchman, L.S. and J.S. Weissman, Native elongating transcript sequencing (NET-seq). Curr Protoc Mol Biol, 2012. Chapter 4: p. Unit 4 14 1–17.

35. Nojima, T., T. Gomes, A.R.F. Grosso, H. Kimura, M.J. Dye, S. Dhir, et al., Mammalian NET-Seq Reveals Genome-wide Nascent Transcription Coupled to RNA Processing. Cell, 2015. 161(3): p. 526–540.

36. Kwak, H., N.J. Fuda, L.J. Core, and J.T. Lis, Precise maps of RNA polymerase reveal how promoters direct initiation and pausing. Science, 2013. 339(6122): p. 950–3.

37. Core, L.J., J.J. Waterfall, D.A. Gilchrist, D.C. Fargo, H. Kwak, K. Adelman, et al., Defining the status of RNA polymerase at promoters. Cell Rep, 2012. 2(4): p. 1025–35.

38. Boija, A., D.B. Mahat, A. Zare, P.H. Holmqvist, P. Philip, D.J. Meyers, et al., CBP Regulates Recruitment and Release of Promoter-Proximal RNA Polymerase II. Mol Cell, 2017. 68(3): p. 491–503 e5.

39. Tresini, M., D.O. Warmerdam, P. Kolovos, L. Snijder, M.G. Vrouwe, J.A. Demmers, et al., The core spliceosome as target and effector of non-canonical ATM signalling. Nature, 2015. 523(7558): p. 53–8.

40. Shibata, A., D. Moiani, A.S. Arvai, J. Perry, S.M. Harding, M.M. Genois, et al., DNA double-strand break repair pathway choice is directed by distinct MRE11 nuclease activities. Mol Cell, 2014. 53(1): p. 7–18.

41. Nielsen, S., Y. Yuzenkova, and N. Zenkin, Mechanism of eukaryotic RNA polymerase III transcription termination. Science, 2013. 340(6140): p. 1577–80.

42. SenGupta, D.J., B. Zhang, B. Kraemer, P. Pochart, S. Fields, and M. Wickens, A three-hybrid system to detect RNA-protein interactions in vivo. Proc Natl Acad Sci U S A, 1996. 93(16): p. 8496–501.

43. Ouyang, J., T. Yadav, J.M. Zhang, H. Yang, E. Rheinbay, H. Guo, et al., RNA transcripts stimulate homologous recombination by forming DR-loops. Nature, 2021.

44. Bottcher, R., M. Hollmann, K. Merk, V. Nitschko, C. Obermaier, J. Philippou-Massier, et al., Efficient chromosomal gene modification with CRISPR/cas9 and PCR-based homologous recombination donors in cultured Drosophila cells. Nucleic acids research, 2014.

45. Kunzelmann, S., R. Bottcher, I. Schmidts, and K. Forstemann, A Comprehensive Toolbox for Genome Editing in Cultured Drosophila melanogaster Cells. G3 (Bethesda, Md), 2016. 6(6): p. 1777–85.

46. Elmer, K., S. Helfer, M. Mirkovic-Hosle, and K. Forstemann, Analysis of endo-siRNAs in Drosophila. Methods in molecular biology, 2014. 1173: p. 33–49.

47. Langmead, B., C. Trapnell, M. Pop, and S.L. Salzberg, Ultrafast and memory-efficient alignment of short DNA sequences to the human genome. Genome Biol, 2009. 10(3): p. R25.

48. Quinlan, A.R., BEDTools: The Swiss-Army Tool for Genome Feature Analysis. Curr Protoc Bioinformatics, 2014. 47: p. 11 12 1–34.

49. Robinson, J.T., H. Thorvaldsdottir, W. Winckler, M. Guttman, E.S. Lander, G. Getz, et al., Integrative genomics viewer. Nat Biotechnol, 2011. 29(1): p. 24–6.

